# Priming is dispensable for NLRP3 inflammasome activation in human monocytes

**DOI:** 10.1101/2020.01.30.925248

**Authors:** Anna Gritsenko, Shi Yu, Fatima Martin-Sanchez, Ines Diaz del Olmo, Eva-Maria Nichols, Daniel M. Davis, David Brough, Gloria Lopez-Castejon

## Abstract

Interleukin (IL)-1 family of cytokines modulate immune responses during infection and inflammation. IL-18 and IL-1β are members of the IL-1 family, which contribute to inflammatory conditions such as rheumatoid arthritis and Alzheimer’s disease. IL-18 and IL-1β are produced as inactive precursors that are activated by large macromolecular complexes called inflammasomes upon sensing damage or pathogenic signals. Canonical NLRP3 inflammasome activation is regarded to require a priming step that causes NLRP3 and IL-1β gene upregulation, and also NLRP3 post-translational licencing. A subsequent activation step leads to the assembly of the inflammasome and the cleavage of pro-IL-18 and pro-IL-1β by caspase-1 into their mature forms, allowing their release. Here we show that in primary human monocytes, the initial priming step is dispensable to form an active NLRP3 inflammasome. We found that, in the absence of priming, the NLRP3 activator nigericin caused the processing and release of constitutively expressed IL-18. Another IL-1 family member, IL-37, is constitutively cleaved but the release of its mature form was mediated by inflammasome activation, also in the absence of a priming step. This NLRP3 activation was characterised by ASC oligomerisation as well as caspase-1 and GSDMD cleavage and was blocked by the NLRP3 inhibitor MCC950 and in NLRP3 deficient cells. IL-18 and IL-37 release were impaired in GSDMD deficient THP-1s, suggesting that pyroptosis is required for release of these cytokines. This work highlights the readiness of the NLRP3 inflammasome to assemble in the absence of priming and hence contribute to sterile inflammatory processes in health and disease.

**Significance Statement:** The NLRP3 inflammasome is a driver of inflammation through the processing of Interleukins (IL)-1β and IL-18. Human monocytes coordinate the innate immune response through inflammasome activation following exposure to pathogens and damage signals. We currently think of NLRP3 activation as a 2 step process: priming (NLRP3 gene upregulation and post-translational licencing) and assembly. Here we show that the priming step is dispensable for NLRP3 inflammasome activation in human monocytes. The second signal alone is sufficient for caspase-1 activation, leading to cell death and the release of the constitutively expressed IL-18 and mIL-37. This reveals that in human monocytes, the NLRP3 inflammasome is already licenced and able to quickly assemble to mount an inflammatory response.

## INTRODUCTION

The Interleukin (IL-1) family plays a vital role in orchestrating inflammatory responses and is involved in many non-communicable diseases such as rheumatoid arthritis, Alzheimer’s disease and cancer [1]. The IL-1 family is comprised of both pro-inflammatory members such as IL-18, IL-1α and IL-1β, as well as anti-inflammatory members that help dampen and control inflammation e.g. IL-1Ra and IL-37 [2]. Cleavage of precursors pro-IL-1β and pro-IL-18 by caspase-1 occurs upon assembly of macromolecular complexes called inflammasomes in response to a variety of danger and pathogenic signals [3], and is required for their activation and release.

NOD-like receptor pyrin domain-containing protein 3 (NLRP3) is the best studied inflammasome sensor. The canonical NLRP3 inflammasome activation is believed to be a two-step process. The first or priming step is achieved through the activation of the nuclear factor kappa B (NF-κB) pathway, leading to the upregulation of NLRP3 and pro-IL-1β proteins [4]. This can be initiated by pathogen-associated molecular patterns (PAMPs) or damage-associated molecular patterns (DAMPs) binding toll like receptors (TLRs), or IL-1β and TNF-α binding to their respective receptors [5]. The second or activating step leads to NLRP3 inflammasome oligomerisation and can be induced by a broad range of factors including PAMPs and DAMPs e.g. nigericin toxin, extracellular ATP, as well as lysosomal destabilisation agents such as silica and cholesterol crystals [6]. Upon inflammasome activation, oligomerised NLRP3 polymerises the adaptor protein ASC, recruiting pro-caspase-1, which undergoes proximity-dependent auto-activation and as a result cleaves pro-IL-18 and pro-IL-1β into their mature forms. Simultaneously, mature caspase-1 induces the cleavage of gasdermin-D (GSDMD) [7] into N-terminal fragments that form lytic pores, facilitating the release of mature IL-1β and IL-18 as well as promoting pyroptotic cell death [8]. GSDMD dependent lytic cell death also leads to the release of damage associated molecules such as HMGB1 (High mobility group box 1), ATP, DNA and even inflammasome components themselves e.g. ASC, which are able to propagate inflammation [9].

Various studies have emerged to suggest that priming is not simply the upregulation of NLRP3 and pro-IL-1β mRNA, but can also involve post-translational modifications (PTM) such as deubiquitination and phosphorylation that licence NLRP3 and promote inflammasome assembly. Several deubiquitinases such as USP7, USP47 and BRCC3 have been reported to licence NLRP3 by promoting its deubiquitination [10, 11]. Changes in NLRP3 phosphorylation by either serine/threonine kinases such as JNK (c-Jun N-terminal kinase) [12], PKA and PKD (protein kinase A and D) [13, 14] or phosphatases such as PP2A (protein phosphatase 2A) [15] and PTPN22 (protein tyrosine phosphatase non-receptor type 22) [16] also modulate inflammasome activation. However whether these contribute to the NLRP3 priming or activating step is not clear.

Whilst pro-IL-1β is not expressed basally and must be induced, pro-IL-18 has been found to be constitutively expressed in macrophages, monocytes, dendritic cells, astrocytes and microglia [17]. IL-37 is another member of the IL-1 family of cytokines that is constitutively expressed. IL-37 has not been widely studied, perhaps because its mouse homologue is still unidentified [18]. Its expression is relatively low in human cells under physiological conditions, but monocytes are the main producers of IL-37 within human peripheral blood mononuclear cells (PBMCs) [19]. Both IL-37 precursor (pro-IL-37) and processed forms (mIL-37) are biologically active, although the mIL-37 monomer has higher anti-inflammatory potency [20]. Pro-IL-37 is thought to be cleaved by caspase-1 and exit the cell via an uncharacterised mechanism following lipopolysaccharide (LPS) stimulation [21] although the role of the inflammasome in this process is not clear.

The current dogma of NLRP3 inflammasome activation is that it requires the presence of a TLR activating ligand. It is accepted that NLRP3 priming is important for the formation of an active inflammasome [22]. However, there is some evidence suggesting that IL-18 is released from human THP-1 cells and human monocytes in the absence of priming [23]. Whether this process is dependent on the NLRP3 inflammasome has not been previously explored. Here, we report that primary human monocytes are equipped with a ready to assemble NLRP3 platform that processes constitutively expressed caspase-1 substrates in a sterile setting. This pathway is dependent on NLRP3, ion efflux and involves post-translational modifications, which are characteristic of the classical primed inflammasome activation. The consequent pyroptotic cell death leads to cytokine processing and release that may subsequently propagate inflammation.

## RESULTS

### Priming is not required for NLRP3 inflammasome activation in human monocytes

To determine if NLRP3 inflammasome activation occurred in human monocytes in the absence of priming, we first compared the release of mIL-18 and mIL-1β, as pro-IL-18 gene expression is constitutive, whilst pro-IL-1β requires upregulation by priming [24]. Undifferentiated THP-1 cells were primed with LPS (1 µg/ml) for 4 hours or left unprimed followed by treatment with the NLRP3 activator nigericin (10 µM, 45 min). We found that nigericin induced mIL-18 release and cell death in both primed and unprimed cells to the same extent, while it only induced mIL-1β release in LPS primed THP-1 cells (Fig 1A, B). In line with this, we detected caspase-1 cleavage into its active form (p20) in both primed and unprimed cells, as well as IL-18 processing and release (Fig 1B). As little is known about the regulation of IL-37 by the inflammasome and it is also constitutively expressed, we investigated the effect of inflammasome activation on IL-37 processing and release under these conditions. Similar to IL-18, western blotting detected two IL-37 fragments in THP-1 supernatants, one of 26 kDa and another of 17.5 kDa, corresponding to pro and mature IL-37 [21] (Fig 1B). These two forms of IL-37 were already present inside the cell in resting conditions indicating that unlike IL-1β and IL-18, IL-37 was cleaved upon expression (Fig 1B). Nigericin treatment, with or without priming, led to GSDMD cleavage, detected by the appearance of a GSDMD fragment of 31 kDa, corresponding to the pore-forming N-terminus (GSDMD NT) (Fig 1B). GSDMD cleavage is a consequence of caspase-1 activation and required for pyroptosis [7]. However, we could not detect significantly elevated levels of cell death following nigericin treatment (Fig 1A). Caspase-1 activation leads to the direct or indirect processing of substrates other than IL-1β and IL-18 [25-27]. For example, capase-1 can mediate the degradation of the E2 ubiquitin-conjugating enzyme UBE2L3 involved in NF-kB activation and pro-IL-1β turnover [28]. To asses if active caspase-1 in unprimed cells can be involved in other cellular processes we investigated whether the second signal alone was sufficient to induce degradation of UBE2L3. Like LPS and nigericin treatment, nigericin treatment alone caused a reduction in UBE2L3 levels, suggesting that caspase-1 is active and able to cleave substrates other than IL-1β or IL-18 (Fig 1B). The NLRP3 protein could be detected in THP-1 cells without any treatment, and is upregulated following exposure to LPS (Fig 1B). Despite this, caspase-1 and GSDMD cleavage, UBE2L3 degradation and mIL-18 and mIL-37 release are not potentiated by LPS stimulation in THP-1 cells. Overall, this highlights that inflammasome activation without prior priming can lead to the cleavage of a number of constitutively expressed caspase-1 substrates.

**Figure 1:**
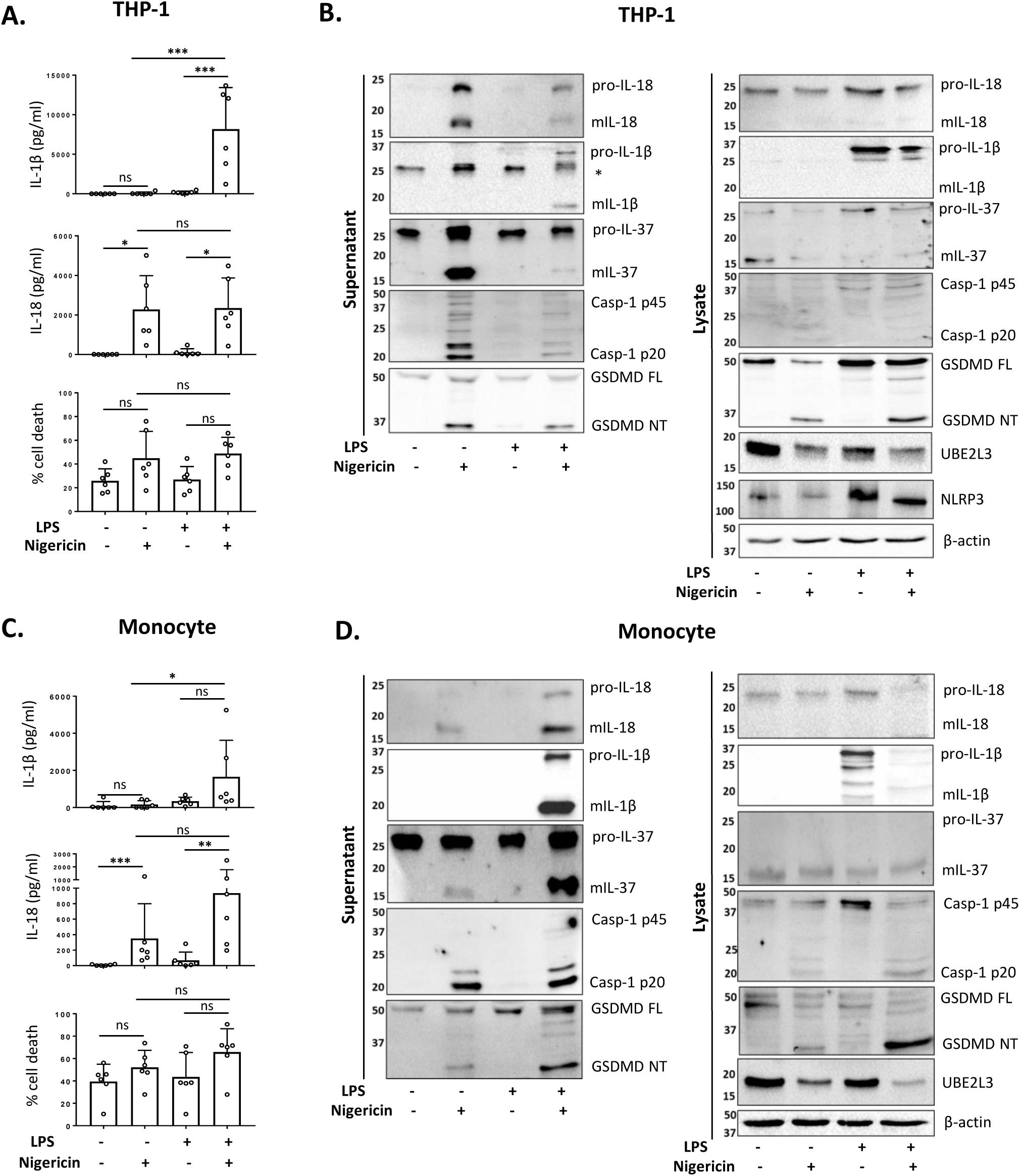
Priming is not required for NLRP3 inflammasome activation in human monocytes. (A, B) Undifferentiated THP-1 cells and (C, D) primary CD14^+^ monocytes were left untreated or primed with LPS (1 µg/mL, 4h) prior to treatment with nigericin (10 µM, 45min) to activate the NLRP3 inflammasome. (A, C) IL-1β and IL-18 were measured by ELISA and cell death was measured by LDH assay and shown as percentage relative to total cell death, n=6. * = P < 0.05; ** = P < 0.01; *** = P <0.001. (B, D) Western blot analysis for mIL-18 (18 kDa), pro-IL-18 (24 kDa), mIL-1β (17 kDa), pro-IL-1β (34 kDa), mIL-37 (17.5 kDa), pro-IL-37 (26 kDa), mCaspase-1 (20 kDa), pro-Caspase-1 (45 kDa), GSDMD full length (FL, 53 kDa), GSDMD N-terminus (NT, 31 kDa), UBE2L3 (17.9 kDa), NLRP3 (113 kDa), as well as loading control β-actin (42 kDa). Blots are representative of at least 3 independent experiments.

Next we tested whether unprimed inflammasome activation occurred in freshly isolated primary human CD14^+^ monocytes from healthy donors. To confirm that the isolated monocytes were not primed at baseline, we cultured them for 4 hours in presence or absence of LPS (1 µg/ml). IL-6 and TNF-α release was used as a measure of priming as these are upregulated and released in response of various signalling pathways such as NF-κB [29]. We found minimal levels of IL-6 and TNF-α in the supernatants of untreated cells, suggesting that the isolated monocytes are unprimed without LPS treatment (Fig 1SA,B). To further confirm that NF-κB signalling is not engaged at baseline, we tested the cell lysates for the presence of inhibitor of kappa B alpha (IκBα), which needs to be phosphorylated and degraded in order for NF-κB for translocate to the nucleus [30]. In untreated primary human monocytes, we detected abundant levels of total IκBα, which were absent when cells were treated with LPS (Fig 1SC). This suggests that NF-κB is sequestered in the cytoplasm and does not drive inflammatory gene expression.

We observed that the treatment of CD14^+^ human monocytes with nigericin alone was sufficient to trigger significant mIL-18 secretion (Fig 1C, D). Western blot analysis revealed that, like THP-1 cells, unprimed monocytes responded to nigericin treatment with caspase-1 activation, GSDMD cleavage, UBE2L3 degradation and the release of mIL-18, mIL-37, but not IL-1β (Fig 1C, D). We observed that priming human monocytes with LPS, unlike THP-1 cells, potentiated nigericin-induced secretion of mIL-18 and cell death (Fig 1C). We have previously seen a very modest upregulation of pro-IL-18 protein following a 4 hour LPS treatment of primary human monocytes, suggesting that LPS potentiates mIL-18 release by acting on the inflammasome. All these data suggested that the priming step is not a requirement for inflammasome activation in human monocytes.

To determine whether unprimed NLRP3 inflammasome assembly is unique to monocytes, we also carried out parallel experiments in human monocyte derived macrophages (MDMs). We observed non-significant mIL-18 release from MDMs in the absence of priming (Fig 2SA). The levels of secreted mIL-18 in this condition were significantly lower than those released by LPS primed cells activated with nigericin. The cleavage of caspase-1 by inflammasome activation without LPS stimulation was also much lower than in LPS primed cells (Fig 2SA). The cleavage of GSDMD and maturation of IL-18 occurred without priming at lower levels than in LPS primed cells (Fig 2SB). The release of mIL-37 was not detected without LPS priming, and UBE2L3 did not appear to be degraded. Therefore, although there is a small response to inflammasome activation in human macrophages, unlike monocytes, priming is required for a full response.

**Figure 2:**
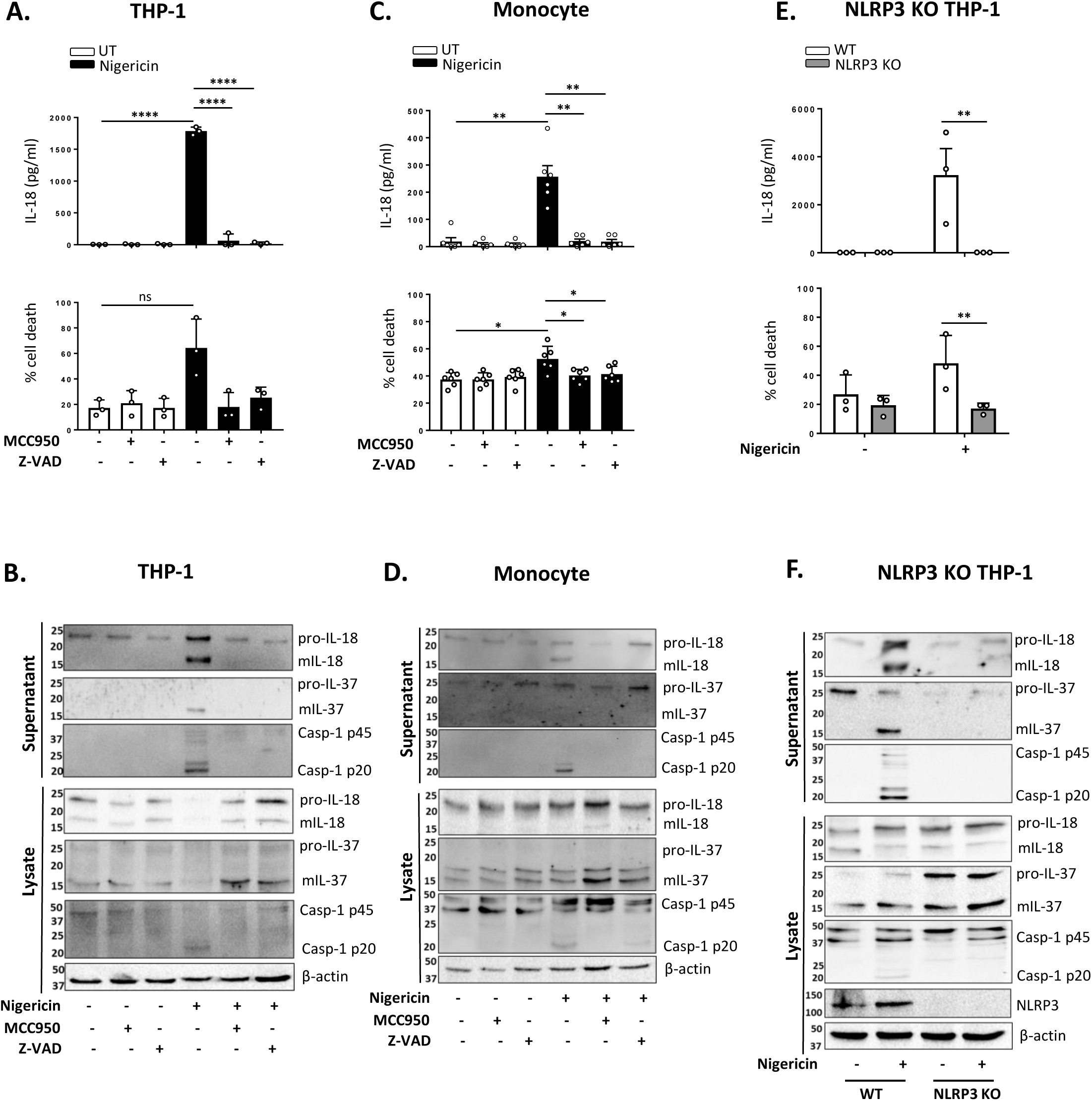
NLRP3 is required for mIL-18 and mIL-37 release in the absence of priming. THP-1 cells (A, B) and monocytes (C, D) were pre-incubated with MCC950 (1 µM) or Z-VAD (50 µM) for 15 minutes prior to treatment with nigericin (10 µM, 45 min). (E, F) Unprimed WT or NLRP3 KO THP-1 were treated with nigericin (10 µM, 45 min). (A, C, E) Secreted IL-18 was measured by ELISA and cell death was measured by LDH assay and shown as percentage relative to total cell death, n=3-6. * = P < 0.05; ** = P < 0.01; *** = P <0.001; **** =P <0.0001. (B, D, F) Western blot analysis for mIL-18 (18 kDa), pro-IL-18 (24 kDa), mIL-37 (17.5 kDa), pro-IL-37 (26 kDa), mCaspase-1 (20 kDa), pro-Caspase-1 (45 kDa), and in some cases NLRP3 (113 kDa), as well as loading control β-actin (42 kDa). Blots are representative of at least 3 independent experiments.

### NLRP3 is required for mIL-18 and mIL-37 release in the absence of priming

To confirm that the mIL-18 and mIL-37 release from unprimed cells is NLRP3 dependent, we initially tested the effect of the NLRP3 inhibitor MCC950 on unprimed inflammasome activation. For this we treated human primary monocytes and THP-1 cells with MCC950 (1 μM, 15 min) prior to the addition of nigericin (10 μM, 45 min) and found that it inhibited nigericin induced mIL-18 release, caspase-1 cleavage into its active form, and cell death in both cell types (Fig 2A-D). Similar effects were observed when either human monocytes or THP-1 cells were pre-treated with an irreversible pan-caspase inhibitor Z-VAD-FMK (ZVAD, 50 μM, 15 min) (Fig 2A-D). Nigericin mediated release of the 17.5 kDa mIL-37 protein was also prevented by MCC950 and Z-VAD in THP-1 cells although both the pro- and mature forms were detected in the cell lysates (Fig 2B), suggesting that the NLRP3 inflammasome mediates mIL-37 release but not pro-IL-37 processing. We observed a high variation in the levels of IL-37 in primary human monocytes in the absence of priming and it could not always be detected, perhaps due to low expression or low stability of the protein in the absence of priming (Fig 2D).

To further confirm the involvement of NLRP3 in this process, we utilised NLRP3 knock out (KO) THP-1 cells and revealed that IL-18 processing and release, caspase-1 activation and cell death following unprimed inflammasome activation did not occur in NLRP3 KO cells compared to WT (Fig 2E-F). Whilst both pro- and mIL-37 were observed in the lysates of WT and NLRP3 KO THP-1s, their release into the supernatant was absent from NLRP3 KO cells treated with nigericin (Fig 2F).

### Unprimed NLRP3 inflammasome activation leads to ASC oligomerisation

NLRP3 inflammasome activation can occur through different pathways [31]. In primed monocytes, nigericin is a trigger of canonical NLRP3 inflammasome activation which is characterised by the oligomerisation of ASC [32]. We therefore tested whether ASC oligomerisation also occurred in unprimed monocytes. We found that nigericin treatment (10 μM, 2 hours) of unprimed THP-1 cells resulted in ASC oligomerisation, detected in the crosslinked NP-40 insoluble fraction of the total lysate, and that this was absent in NLRP3 KO cells (Fig 3A). Similarly, unprimed primary human monocytes formed oligomers following stimulation with nigericin and this did not occur in monocytes pre-treated with the NLRP3 inhibitor MCC950 (10 μM, 15 min) (Fig 3B).

**Figure 3:**
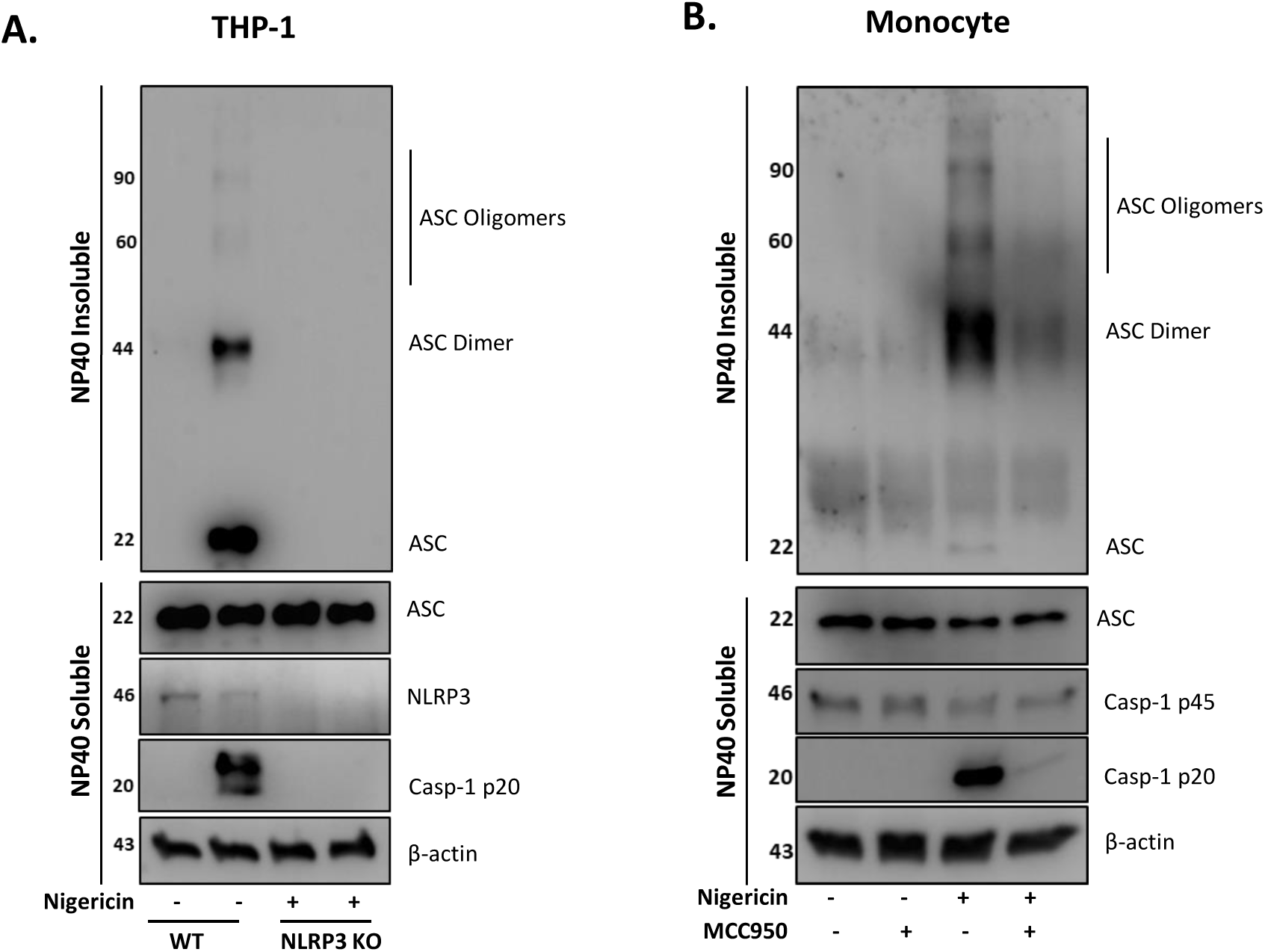
Unprimed NLRP3 inflammasome activation leads to ASC oligomerisation. (A) WT and NLRP3 KO THP-1 cells were left unprimed prior to treatment with nigericin (10 µM, 2h) to activate NLRP3 inflammasome. (B) Monocytes were pre-incubated with MCC950 (1 µM) for 15 minutes prior to treatment with nigericin (10 µM, 45min). The NP-40 soluble fraction and DSS crosslinked insoluble fraction were immunoblotted for ASC monomers, dimers and oligomers. Blots are representative of at least 3 independent experiments.

### Unprimed NLRP3 inflammasome activation has the same requirements as the canonical inflammasome

As well as ASC oligomerisation, the classical primed canonical NLRP3 inflammasome activation is dependent on potassium (K^+^) [33] and chloride (Cl^-^) [34] efflux. This is unlike the alternative NLRP3 inflammasome that only occurs in human monocytes and that assembles following prolonged priming signals in a K^+^ independent manner [34]. We tested the requirement of K^+^ and Cl^-^ efflux during unprimed inflammasome activation. We incubated unprimed THP-1 cells either in high K^+^ containing media or pre-treated these cells with broad-spectrum chloride channel inhibitor NPPB (50 μM) [35], or the NLRP3 inhibitor NBC19 (25 μM) [36] for 15 minutes prior to nigericin treatment (Fig 4A,B). This revealed that unprimed mIL-18 release, caspase-1 cleavage and cell death following nigericin stimulation were dependent on K^+^ efflux and were also sensitive to Cl^-^ channel inhibitor NPPB (Fig 4A,B) as they were reduced to the same extent as in the presence of inflammasome inhibitor NBC19. Moreover, the release of mIL-37 was also blocked in these conditions (Fig 4B).

**Figure 4:**
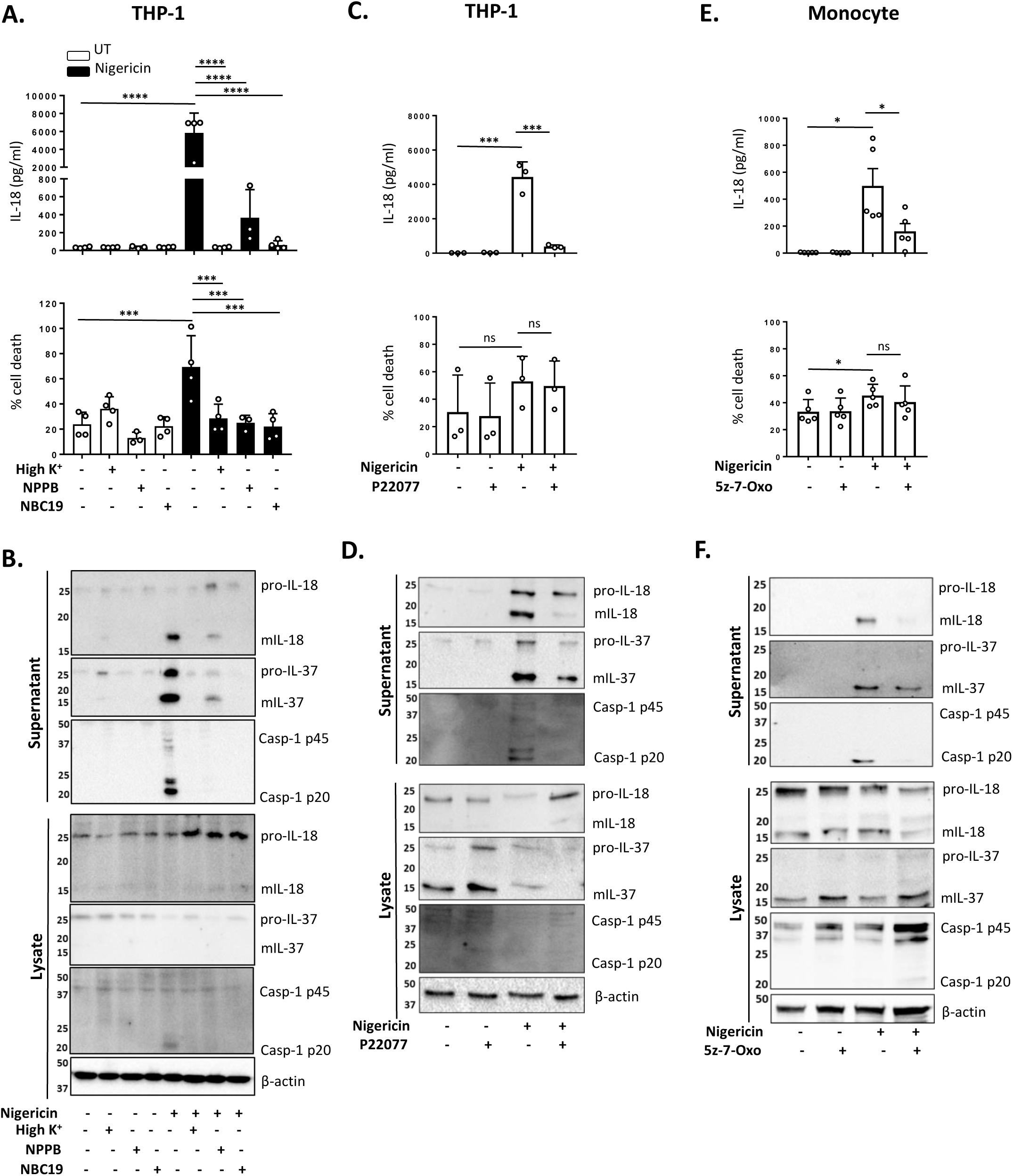
Unprimed NLRP3 inflammasome activation has the same requirements as the canonical inflammasome. (A, B) THP-1 cells were left unprimed and pre-incubated with high K^+^ buffer, NPPB (50 µM) or NBC19 (25 µM) for 15 minutes prior to treatment with nigericin (10 µM, 45min). (C, D) THP-1 cells were left unprimed pre-incubated with P22077 (2.5 µM, 15 min) prior to treatment with nigericin (10 µM, 45 min). (E, F) Primary human monocytes were pre-incubated with 5z-7-Oxozeaenol (0.5 µM) for 15 minutes prior to treatment with nigericin (10 µM, 45 min). (A, C, E) Secreted IL-18 was measured by ELISA and cell death was measured by LDH assay and shown as percentage relative to total cell death, n=3-5. * = P < 0.05; ** = P < 0.01; *** = P <0.001; **** =P <0.0001. (B, D, F) Supernatants and lysates were analysed for mIL-18 (18 kDa), pro-IL-18 (24 kDa), mIL-37 (17.5 kDa), pro-IL-37 (26 kDa), mCaspase-1 (20 kDa), pro-Caspase-1 (45 kDa), as well as loading control β-actin (42 kDa). Blots are representative of at least 3 independent experiments.

Deubiquitination of NLRP3 by both the priming and activating step have been described as necessary to form an active NLRP3 inflammasome [37]. We have recently shown that deubiquitinases USP7 and USP47 mediate canonical NLRP3 inflammasome activation in human macrophages independently of transcriptional priming [11], suggesting they might also mediate inflammasome activation in unprimed human monocytes. To test this we compared the effect of the dual USP7/USP47 inhibitor P22077 [11] on inflammasome activation without priming. We found that pre-incubation of THP-1 cells with P22077 (2.5 µM, 15 min) prior to nigericin treatment significantly reduced IL-18 release (Fig 4C). This was accompanied by a reduction in caspase-1 maturation and mIL-37 release (Fig 4D) although no significant reduction in cell death was observed. These data suggest that in human monocytes, unprimed NLRP3 inflammasome assembly is mechanistically similar to the canonical primed NLRP3 inflammasome activation.

TAK1 (transforming growth factor beta-activated kinase 1) can mediate canonical NLRP3 inflammasome assembly and ASC oligomerisation independently of its role in the NF-kB pathway (inflammasome priming signal) [38]. It has been recently suggested that in human cells, the inflammasome priming step allows for NLRP3 post-translational modifications that lead to cells bypassing the requirement for NEK7, thought to be a critical co-factor for canonical NLRP3 activation [39], in a TAK1 dependent manner [40]. Therefore, we investigated whether TAK1 activity is a requirement for NLRP3 activation in the absence of priming in human monocytes. To assess this, we pre-treated cells with an inhibitor of TAK1 catalytic activity: 5z-7-oxozeaenol (5z-7) (0.5 µM) for 15 minutes prior to nigericin stimulation. We found that TAK1 inhibitor significantly impaired mIL-18 release in primary human monocytes (Fig 4E,F) as well as reduced the release of mIL-37 and the cleavage of caspase-1 into its mature form (p20) (Fig 4F). This indicates that indeed TAK1 plays a role in inflammasome activation in a transcription independent manner.

### mIL-18 and mIL-37 release from unprimed monocytes is dependent on GSDMD

NLRP3 inflammasome mediated IL-1 release and pyroptotic cell death are closely associated, and these events are tightly linked to GSDMD cleavage and pore formation. However, certain NLRP3 activators such as changes in cell volume can induce mIL-1β and mIL-18 release in the absence of pyroptosis [41]. To determine whether inflammasome-induced cell death in unprimed monocytes was involved in the release of mIL-18 and mIL-37, we tested the effect of the polyphenolic compound punicalagin. Punicalagin inhibits plasma membrane permeability, as well as mIL-1β and mIL-18 release induced by inflammasome, whilst allowing caspase-1 activation [42, 43]. In unprimed THP-1 cells, 15 minute pre-treatment with 50 µM punicalagin prior to nigericin significantly blocked IL-18 release (Fig 5A). Western blot analysis revealed that punicalagin treated cells still underwent caspase-1 activation, indicating inflammasome activation, but the release of mature caspase-1 (p20) was blocked (Fig 5B). Similarly, pro-IL-18 was cleaved into its mature form but its release was inhibited by punicalagin. Release of mIL-37 was also prevented by punicalagin treatment, suggesting its release was dependent on membrane permeabilisation (Fig 5B).

**Figure 5:**
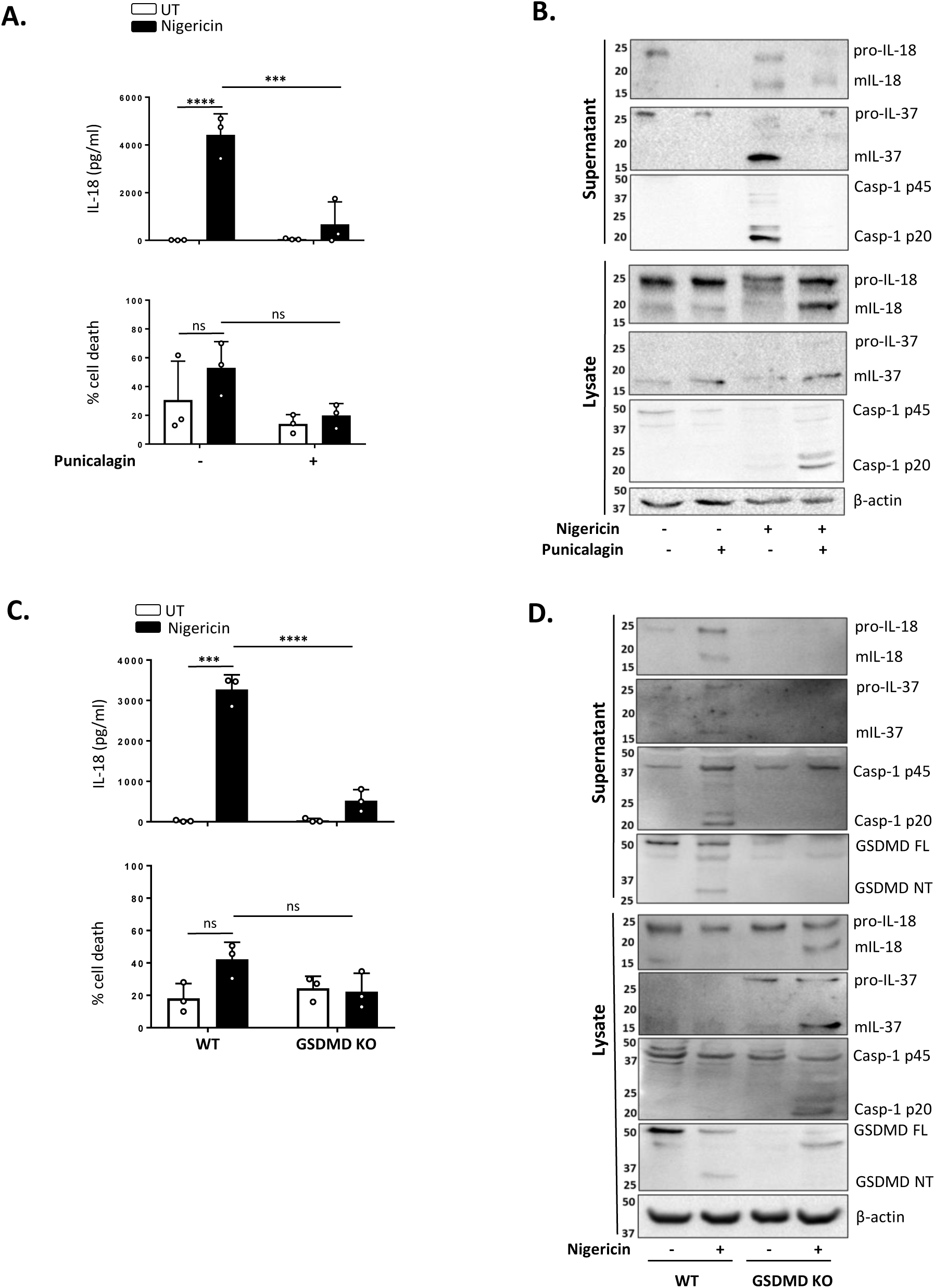
mIL-18 and mIL-37 release from unprimed monocytes is dependent on GSDMD. (A, B) Unprimed THP-1 cells pre-incubated with punicalagin (50 µM, 15 min) prior to treatment with nigericin (10 µM, 45 min). (C, D) Unprimed WT and GSDMD KO THP-1s were treated with nigericin (10 µM, 45 min). (A, C) Secreted IL-18 was measured by ELISA and cell death was measured by LDH assay and shown as percentage relative to total cell death n=3. *** = P <0.001; **** =P <0.0001. (B, D) Western blot analysis of THP-1 cells for mIL-18 (18 kDa), pro-IL-18 (24 kDa), mIL-37 (17.5 kDa), pro-IL-37 (26 kDa), mCaspase-1 (20 kDa), pro-Caspase-1 (45 kDa), GSDMD full length (FL, 53 kDa), GSDMD N-terminus (NT, 31 kDa), as well as loading control β-actin (42 kDa). Blots are representative of at least 3 independent experiments.

The effect of punicalagin on inflammasome activation resembles that of GSDMD KO [42-44]. Similar to pro-IL-18, GSDMD is constitutively expressed in unprimed THP-1 and human monocytes. As the GSDMD pore has been proposed as a mediator of mIL-1β release [44] and we observed GSDMD cleavage in unprimed monocytes, we investigated if GSDMD was required for mIL-18 and mIL-37 release. IL-18 release caused by nigericin was significantly lower in cells lacking GSDMD compared to WT control (Fig 5C). We also found that release of both pro and mature forms of IL-37 was reduced in GSDMD KO THP-1 cells (Fig 5D), suggesting that GSDMD is required for the release of mIL-37 in response to inflammasome activation. Overall, these data suggest that caspase-1 cleavage of GSDMD and subsequent pyroptosis may be a key mechanism for the release of inflammasome regulated cytokines following inflammasome assembly in the absence of priming.

## DISCUSSION

Our work shows that NLRP3 inflammasome activation can occur in the absence of a priming step in human monocytes. The NLRP3 inflammasome priming step has long been regarded as an essential stage for the assembly of an active NLRP3 inflammasome. Initially, it was believed that priming was only required to upregulate NLRP3 and pro-IL-1β proteins in response to the activation of NF-kB signalling [4]. In recent years, however, a role for priming independent of transcriptional regulation has been uncovered and is believed to be mediated by post translational modifications such as changes in the ubiquitination or phosphorylation of NLRP3 [37] [15]. We now show that this initial priming step is not required for the formation of an active inflammasome but may enhance inflammasome activation.

There is evidence linking the ability of human monocytes to activate caspase-1 to the processing and release IL-18 in the absence of priming. Mehta *et al*. (2000) initially showed that human monocytes can activate caspase-1 in response to ATP mediated P2×7R activation in the absence of priming [45]. It was later shown that lysosomal destabilization induced by nigericin contributes to caspase-1 activation and IL-18 relase in what was thought to be necrosis in THP-1 cells and human monocytes [23]. Our data show that this activation of caspase-1 and release of IL-18 in unprimed human monocytes and THP-1 cells is dependent on the activation of the canonical NLRP3 inflammasome as it leads to the formation of ASC oligomers and is a process blocked by preventing K^+^ efflux and Cl^-^ channel inhibition.

It is now evident that changes in post-translational modifications such as ubiquitination or phosphorylation are essential for the assembly of an active inflammasome by arranging NLRP3 into a licenced state [37] [15]. Deubiquitination of NLRP3 is an important step both at the priming and activating level [37] [11]. We have recently found that deubiquitinases USP7 and USP47 are required for canonical NLRP3 inflammasome activation independently of transcriptional priming [11]. We also found that nigericin treatment induced an increase in USP7 and USP47 activity in primed and unprimed macrophages [11], suggesting that USP7 and USP47 could act as direct sensors of danger signals. Here we reveal that blocking their deubiquitinase activity with the inhibitor P22077 prevents the canonical NLRP3 inflammasome activation in unprimed monocytes. Our work highlights the role of USP7 and USP47 in sensing danger and facilitating a fast inflammatory response via unprimed inflammasome assembly.

TAK1 kinase is described to regulate the canonical NLRP3 inflammasome at both priming and activation stages, by either mediating transcriptional priming through its role in NF-κB signalling downstream of TLR4 [46] or by controlling lysosomal rupture that leads to NLRP3 assembly and activation [38]. More recently, TAK1 has been suggested to mediate inflammasome activation via non transcriptional priming [40]. It was reported that the requirement for the NLRP3-binding protein NEK7 to form and active NLRP3 inflammasome [39] can be bypassed via TAK1 mediated potential post translational priming of NLRP3 [40]. It is therefore not clear whether TAK1 plays a role in a setting where TLR4 is not engaged. In unprimed primary human monocytes, we found that TAK1 inhibition significantly reduces inflammasome activation. This shows that independently of its role in the priming step, the catalytic activity of TAK1 is important for NLRP3 licencing and inflammasome assembly induced by the activating signal. However, the downstream target of TAK1 that post translationally licences NLRP3 still remains to be identified.

In addition to mIL-18, we found that mIL-37 is released in response to NLRP3 inflammasome activation in the absence of priming. Although two pro-IL-37 cleavage sites have been described, the contribution of the inflammasome, caspase-1 activation and cell death to its processing and release remains unclear. The first described site, Val46, was identified in COS7 cells overexpressing IL-37. However, the protease responsible for this cleavage remains unknown [47]. The second, Asp20, is a caspase-1 predicted cleavage site. It was reported that mouse RAW cells expressing human pro-IL-37 D20A mutant had reduced pro-IL-37 cleavage upon LPS stimulation compared with wild type pro-IL-37 [21]. However, whether pro-IL-37 cleavage and release could be blocked by caspase-1 inhibition was not explored. There is also some evidence to suggest that IL-37 is secreted via the classical ER-Golgi pathway in response to LPS but whether this was the precursor or the cleaved form was not shown [19]. Our data show that mIL-37 is constitutively present in human monocytes and THP-1 cells and that this processing is not caspase-1 dependent as treatment with Z-VAD or NLRP3 inhibitor MCC950 does not prevent it. We also showed that in the absence of priming, mIL-37 release, like IL-18, is mediated by pyroptosis since treatment of monocytes with punicalagin or the absence of the pore forming protein GSDMD prevented their release but not their processing. These findings provide new evidence for the role of the inflammasome in the release of mIL-37.

Activation of caspase-1 by the NLRP3 inflammasome leads to the processing of IL-1 family members IL-1β and IL-18 [3]. In LPS primed cells it also leads to the degradation of UBE2L3 (E2 ubiquitin-conjugating enzyme L3), involved in pro-IL-1β turnover [28]. We now know that UBE2L3 degradation also occur in an unprimed setting indicating the ability of active caspase-1 to cleave same substrates independently of priming and suggesting that UBE2L3 might be important for other outcomes of inflammasome activation. Our work also reveals the ability of unprimed monocytes to cleave GSDMD and cause pyroptotic cell death. Pyroptosis facilitates the release of various alarmins, such as HMGB1 [9] that could contribute to the amplification of the inflammatory response initiated by IL-18 in an unprimed setting. This however remains unexplored and needs to be further researched.

Pro-IL-18, unlike pro-IL-1β, is constitutively expressed and plays important roles in pathological conditions where no priming signals are involved. This is especially relevant in autoinflammatory syndromes caused by mutations of NLRP3 [48], where IL-18R deletion in a cryopyrin-associated periodic syndromes (CAPS) mouse model lessened the disease phenotype to a greater extent than IL-1R deletion [49]. Pyrin mutations associated with familial Mediterranean fever (FMF) [50] and the NLRC4 gain of function mutations that mediate the Macrophage Activation Syndrome (MAS) [51] are all characterised by systemically elevated IL-18 levels. Moreover, the release of damage associated signals such as ATP in transplantation can lead to P2×7R-mediated activation of the inflammasome inducing the release of IL-18, but not IL-1β, which in turn plays an important role in allograft rejection through IFN-γ production and CD8^+^ T cell proliferation [52]. Inflammasomes are also activated during the progression of sterile inflammatory diseases including atherosclerosis, metabolic and neuro-inflammatory disorders where IL-18 is an important driver [53]. In these conditions, IL-18 mediates the activation of pro-inflammatory NF-κB and upregulation and secretion of TNF-α, adhesion molecules and chemokines [54]. This highlights the importance of the ability of the NLRP3 inflammasome to rapidly assemble without the need to transcriptionally upregulate its components, especially in monocytes.

It is appreciated that inflammatory conditions such as Sjögren’s autoimmune disease are characterised by increased levels of IL-37, as well as IL-18 [55]. Therefore, the parallel release of IL-18 and IL-37 by human monocytes described here could respond to a need to control and limit inflammatory damage during disease. The dynamics of the release and activity of these cytokines during inflammation are poorly understood and more research in this area is required specially when considering the use of IL-37 as potential anti-inflammatory therapy [56].

Based on our results, we propose that the NLRP3 inflammasome in human cells is a ‘ready to assemble’ complex, capable of propagating IL-18 or IL-37 mediated responses in the absence of priming and is therefore essential in the initial steps of inflammatory pathways both in health and disease.

## MATERIALS AND METHODS

### Reagents and Antibodies

LPS (*Escherichia coli* 026:B6); nigericin sodium salt (N7143); protease inhibitor cocktail (P8340); penicillin–streptomycin (Pen/Strep, P4333); MCC950 (PZ0280); Punicalagin (P0023), 5Z-7-Oxozeaenol (O9890) and Z-VAD-FMK (V116-2MG) were obtained from Sigma. Foetal bovine serum (FBS) was purchased from Gibco. 5-Nitro-2-(3-phenylpropylamino)benzoic acid (NPPB, 0593) was sourced from Calbiochem.

Primary antibodies for Western blot assays and their final concentrations were as follows: anti-human IL-18 (0.5 μg/ml, rabbit polyclonal, LifeSpan BioSciences, LS-C313397), anti-human IL-1β (0.1 μg/ml, goat polyclonal, R&D Systems, AF-201-NA), anti-human IL-37 (2 μg/ml, goat polyclonal, R&D Systems, AF1975), anti-human caspase-1 p20 (1:1000, mouse monoclonal, Cell Signalling Technology, 3866), anti-NLRP3 (1 μg/ml, mouse monoclonal, Adipogen, AG-20B-0014), and anti-β-actin-HRP (0.2 μg/ml, mouse monoclonal, Sigma, A3854), anti-human UBE2L3 (0.2 μg/ml, mouse monoclonal, Santa Cruz, sc-390032), anti-human GSDMD (0.07 μg/ml, Novus Biologicals, NBP2-33422), anti-human IκBα (1:1000, Cell Signalling Technology 4812). HRP conjugated secondary antibodies used for Western blotting were anti-rabbit-HRP (0.25 μg/ml, goat polyclonal, Dako, P0448), anti-mouse-HRP (1.3 μg/ml, rabbit polyclonal, Dako, P0260) and anti-goat-HRP (0.13 μg/ml, rabbit polyclonal, Sigma, A5420).

### Cell Culture and Treatments

THP-1 cells were cultured in RPMI-1640 supplemented with 2 mM L-glutamine, 10% FBS and Pen/Strep (100 U/ml) and plated at a density of 1 × 10^6^ cells/ml. Human peripheral blood mononuclear cells (PBMCs) were obtained from the National Blood Transfusion Service (Manchester, UK) with full ethical approval from the Research Governance, Ethics, and Integrity Committee at the University of Manchester (ref. 2018-2696-5711). PBMCs were isolated from leucocyte cones from healthy donors by density centrifugation using a 30% Ficoll gradient. The PBMC layer was separated and washed with MACS buffer (PBS, 0.5% BSA, 2mM EDTA) to remove platelets. Monocytes were positively selected from PBMCs with magnetic CD14^+^ MicroBeads (Miltenyi, 130-050-201) for 15 minutes at 4°C and eluted using a LS column (Miltenyi, 130-042-401). To differentiate monocyte-derived macrophages (MDMs), monocytes were plated for 7 days (at a concentration of 5 × 10^5^ cells per millilitre) in RPMI-1640 supplemented with 2 mM L-glutamine, 10% FBS, Pen/Strep (100 U/ml) and 0.5 ng/ml M-CSF (Peprotech, 300-25). On day 3, half of the media was removed and replaced with fresh media.

GSDMD knockout THP-1 cells were lentivirally generated using guide RNA oligonucleotides sequence 5’-CACCGCTGCAAGCTGGCCAGGTACC-3’ and 5’-AAACGGTACCTGGCCAGCTTGCAGC-3’ [57] by utilising the lentiCRISPR v2 plasmid system. lentiCRISPR v2 was a gift from Feng Zhang (Addgene plasmid # 52961 ; http://n2t.net/addgene:52961; RRID:Addgene_52961). NLRP3 deficient THP-1 cells were a gift from Prof Veit Hornung (Ludwig Maximilian University of Munich).

Seeded cells were left unprimed or primed for 4 hours with LPS (1 μg/ml). The priming stimulus was then removed and replaced with ET buffer (147 mM NaCl, 10 mM HEPES, 13 mM D-glucose, 2 mM KCl, 2 mM CaCl_2_ and 1 mM MgCl_2_). Cells were treated Nigericin toxin (10 μM, 45 minutes or 2 hours as indicated) to activate the NLRP3 inflammasome.

### Cell Death Assay

Cell death was measured using quantitative assay for the release of lactate dehydrogenase (LDH) into cell supernatants. The supernatant was gently centrifuged for 5 minutes at 500 ×g at 4°C to remove any remaining cells. LDH release in cell supernatants were measured using CytoTox 96® Non-Radioactive Cytotoxicity Assay (G1780, Promega), according to the manufacturer’s instructions. Absorbance values were recorded at 490 nm and the results were expressed as a percentage of LDH release normalised to total lysis.

### Enzyme-linked immunosorbent assay (ELISA)

Levels of human IL-1β (DY201) and IL-18 (DY318) were measured in the cell supernatants using ELISA kits from R&D Systems. Human IL-6 (# 88-7066-86) and TNF-α (# 88-7346-86) were detected using Invitrogen ELISA kits. ELISAs were performed following the manufacturer’s instructions.

### Western Blot

Cells were lysed on ice using a RIPA lysis buffer (50 mM Tris–HCl, pH 8, 150 mM NaCl, 1% NP-40, 0.5% sodium deoxycholate and 0.1% sodium dodecyl sulphate, SDS), supplemented with a protease inhibitors cocktail (Sigma-Aldrich, P8340, 1:100). Lysates were then clarified at 21,000 ×g for 10 minutes in order to remove the insoluble fraction. Protein concentrations of each sample were measured using BCA assays (Thermo Scientific Pierce, 23225), following the manufacturer’s guidelines, so an equal amount of protein was loaded for each sample. Cell supernatants were centrifuged at 500 ×g for 5 minutes to remove dead cells and concentrated using 10 kDa MW cut-off filters (Amicon, Merck Millipore), as described by the manufacturer. Supernatants and whole-cell lysates were diluted in 1X reducing Laemmli buffer containing 1% β-mercaptoethanol. Samples were boiled at 95°C for 5 minutes and separated by Tris-glycine SDS–PAGE. Proteins were transferred onto nitrocellulose membranes (0.2 µm) and blocked in PBS-Tween (PBS-T, 0.1%) containing 5% skimmed milk for 1 hour at room temperature. Membranes were then incubated overnight with the specific primary antibody in blocking buffer at 4°C. The following day, membranes were labelled with a horseradish peroxidase-conjugated secondary antibody for 1 hour at room temperature. Membranes were then washed, developed and captured digitally using Clarity™ Western ECL Blotting Substrate (Bio-Rad, 1705061) in a ChemiDoc™ MP Imager (Bio-Rad).

### ASC Oligomerisation Assay

After inflammasome stimulation, undifferentiated WT and NLRP3 KO THP-1 cells and monocytes were placed on ice. 1% (v/v) NP-40 and protease inhibitor cocktail were added directly to wells. Cell total lysates were separated into NP-40 soluble and insoluble fractions using differential centrifugation at 6,800×g for 20 minutes at 4°C. The soluble fraction containing cell supernatant and lysates were utilised for western blotting analysis, whereas the NP-40 insoluble pellets were chemically crosslinked with 2 mM disuccinimidyl suberate (DSS) (Thermo Fisher) for 30 minutes at RT. Crosslinked pellets were further centrifuged at 6,800×g for 20 minutes and resuspended in boiled 1x Laemmli buffer for standard SDS-PAGE.

### Statistical Analysis

GraphPad Prism 8 software was used to carry out all statistical analysis. Differences between 2 groups were analysed using t-test. Differences between 3+ groups were analysed using one-way ANOVA with the *post hoc* Dunnett’s test or two-way ANOVA with the *post hoc* Tukey’s test for multiple comparisons. Data was shown as mean +/- standard deviation. Accepted levels of significance were *P < 0.05, **P < 0.01, ***P < 0.001, ****P < 0.0001.

## Acknowledgments

We thank Dr. Veit Hornung (Ludwig Maximilian University, Munich) for providing the NLRP3 KO THP-1 cells and Dr. Kevin Stacey (Lydia Becker Institute of Immunology and Inflammation, Manchester) for his support in providing PBMCs.

## Contributions

A.G., S.Y.,, D.B. and G.L-C. designed research; A.G., S.Y., F. M-S., I. D-O., D.B. and G.L-C. performed research; D.D. and E. M-N. contributed new reagents/analytic tools; A.G., S.Y., F. M-S.,D.B and G.L.-C. analysed and discussed data; A.G. and G. L-C. wrote the paper.

