## Supplemental Fig 1 for "Priming is dispensable for NLRP3 inflammasome activation in human monocytes"

### Slide 1
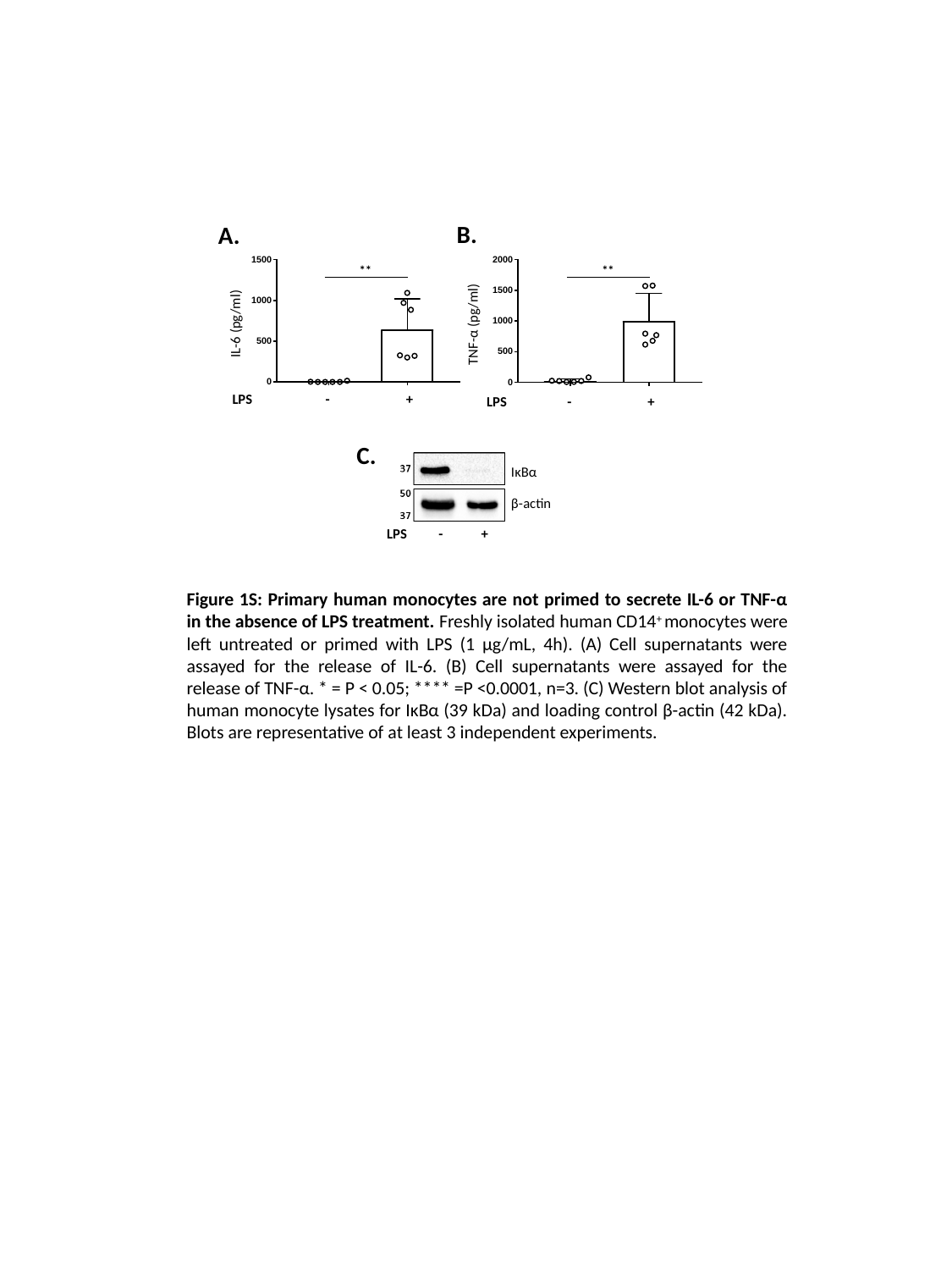

B.
A.
**
**
IL-6 (pg/ml)
TNF-α (pg/ml)
LPS - +
LPS - +
C.
IκBα
β-actin
LPS - +
Figure 1S: Primary human monocytes are not primed to secrete IL-6 or TNF-α in the absence of LPS treatment. Freshly isolated human CD14^+^ monocytes were left untreated or primed with LPS (1 µg/mL, 4h). (A) Cell supernatants were assayed for the release of IL-6. (B) Cell supernatants were assayed for the release of TNF-α. * = P < 0.05; **** =P <0.0001, n=3. (C) Western blot analysis of human monocyte lysates for IκBα (39 kDa) and loading control β-actin (42 kDa). Blots are representative of at least 3 independent experiments.
