## Supplemental Fig 2 for "Priming is dispensable for NLRP3 inflammasome activation in human monocytes"

### Slide 1
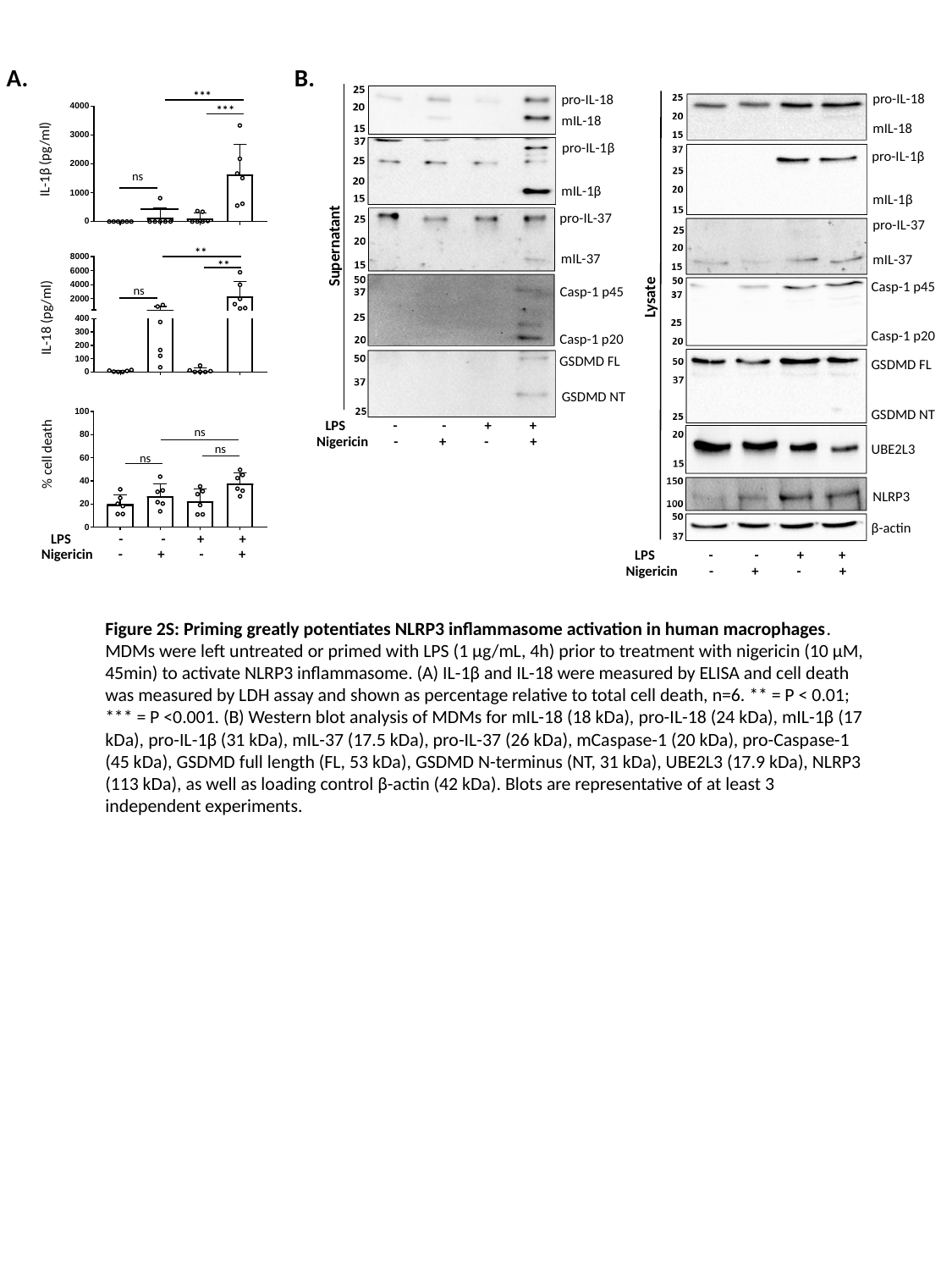

B.
A.
***
pro-IL-18
pro-IL-18
***
mIL-18
mIL-18
pro-IL-1β
pro-IL-1β
IL-1β (pg/ml)
Supernatant
ns
mIL-1β
mIL-1β
pro-IL-37
pro-IL-37
**
mIL-37
mIL-37
**
Lysate
Casp-1 p45
Casp-1 p45
ns
IL-18 (pg/ml)
Casp-1 p20
Casp-1 p20
GSDMD FL
GSDMD FL
GSDMD NT
GSDMD NT
LPS - - + +
ns
Nigericin - + - +
UBE2L3
ns
% cell death
ns
NLRP3
β-actin
LPS - - + +
Nigericin - + - +
LPS - - + +
Nigericin - + - +
Figure 2S: Priming greatly potentiates NLRP3 inflammasome activation in human macrophages. MDMs were left untreated or primed with LPS (1 µg/mL, 4h) prior to treatment with nigericin (10 µM, 45min) to activate NLRP3 inflammasome. (A) IL-1β and IL-18 were measured by ELISA and cell death was measured by LDH assay and shown as percentage relative to total cell death, n=6. ** = P < 0.01; *** = P <0.001. (B) Western blot analysis of MDMs for mIL-18 (18 kDa), pro-IL-18 (24 kDa), mIL-1β (17 kDa), pro-IL-1β (31 kDa), mIL-37 (17.5 kDa), pro-IL-37 (26 kDa), mCaspase-1 (20 kDa), pro-Caspase-1 (45 kDa), GSDMD full length (FL, 53 kDa), GSDMD N-terminus (NT, 31 kDa), UBE2L3 (17.9 kDa), NLRP3 (113 kDa), as well as loading control β-actin (42 kDa). Blots are representative of at least 3 independent experiments.
